# Meiotic Cas9 expression mediates genotype conversion in the male and female mouse germline

**DOI:** 10.1101/2021.03.16.435716

**Authors:** Alexander J. Weitzel, Hannah A. Grunwald, Rimma Levina, Valentino M. Gantz, Stephen M. Hedrick, Ethan Bier, Kimberly L. Cooper

## Abstract

Highly efficient genotype conversion systems have potential to facilitate the study of complex genetic traits using laboratory mice and to limit loss of biodiversity and disease transmission caused by wild rodent populations. We previously showed that such a system of genotype conversion from heterozygous to homozygous after a sequence targeted CRISPR/Cas9 double strand DNA break is feasible in the female mouse germline. In the male germline, however, all double strand breaks were instead repaired by end joining mechanisms to form an ‘insertion/deletion’ (indel) mutation. These observations suggested that timing Cas9 expression to coincide with meiosis I is critical to favor conditions when homologous chromosomes are aligned and interchromosomal homology directed repair (HDR) mechanisms predominate. Here, using a Cas9 knock-in allele at the *Spo11* locus, we show that meiotic expression of Cas9 does indeed mediate genotype conversion in the male as well as in the female germline. However, the low frequency of both HDR and indel mutation in both male and female germlines suggests that Cas9 may be expressed from the *Spo11* locus at levels too low for efficient double strand DNA break formation. We suggest that more robust Cas9 expression initiated during early meiosis I may improve the efficiency of genotype conversion and further increase the rate of ‘super-Mendelian’ inheritance from both male and female mice.

## Introduction

The mouse remains the single most utilized laboratory model of human physiology and disease. Yet, breeding strategies that combine multiple homozygous transgenic or knock-in alleles are cumbersome in mice due to small litter sizes and long generation times relative to other traditional model systems. Practical challenges therefore limit the full potential for mouse genetic research to model certain complex human genetic disorders and to engineer models that better mimic human metabolism and immunity for therapeutic drug design. We previously demonstrated that super-Mendelian inheritance mediated by CRISPR/Cas9 is feasible through the female germline of mice, increasing the transmission frequency of a transgenic allele [1]. Similar systems have been proposed for implementation in wild rodents over multiple generations to limit the spread of infectious disease or to reduce populations of invasive mice and rats in sensitive island ecosystems [2–5].

To assess the feasibility of CRISPR/Cas9-mediated genotype conversion, we previously engineered a ‘CopyCat’ transgene that disrupts the fourth exon of *Tyrosinase*, a gene that is required for melanization in mice [6]. The *Tyr*^*CopyCat*^ transgene expresses the guide RNA (gRNA) used to target the site of its own insertion and an mCherry fluorescent marker to track its inheritance. We then used available genetic tools to express Cas9 in the early embryo and in the male and female germline.

There are two ‘non-conservative’ repair outcomes after CRISPR/Cas9-mediated double strand DNA break (DSB) that alter the sequence of the wild type target site in a *Tyr*^*CopyCat*^ heterozygous animal. First, the two ends may be re-joined by non-homologous end joining or micro-homology mediated end joining [NHEJ or MMEJ, hereafter referred to as ‘end joining’ (EJ)], which can cause insertion or deletion mutations (indels) that disrupt the gRNA target site and prevent subsequent cutting (reviewed in [7]). Alternatively, the DSB can be repaired by homology directed repair (HDR) using genomic sequences that flank the *Tyr*^*CopyCat*^ transgene on the homologous chromosome. If interchromosomal HDR occurs, then the transgene that was inherited on only one chromosome is copied into the DSB of the homologous locus such that its genotype is converted from heterozygosity to homozygosity [1] (Fig 2A).

CRISPR/Cas9-mediated genotype conversion is highly efficient (>90%) in Dipteran insects and in yeast [8–11]. Using multiple Cre/lox strategies to initiate Cas9 expression in oogonia and spermatogonia of mice [12,13], we previously demonstrated genotype conversion in the female but not in the male germline [1]. In females, the frequency of allele transmission increased from 50%, predicted by Mendelian inheritance, to 72% after CRISPR/Cas9-mediated genotype conversion. We also recombined ultra-tightly linked loci (∼9 kb apart) by genotype conversion in 22.5% of offspring of females. Although this was an important demonstration of the powerful potential of such approaches, use of two transgenes (Cre and conditional Cas9) to achieve timing is not an optimal approach to simplify complex crossing schemes.

The observation that genotype conversion occurred exclusively in the female germline suggested that sex differences in the progression of gonial development might be key. Oogonia rapidly mature into primary oocytes that initiate meiosis during fetal development [14], while spermatogonia are mitotic throughout the life of the animal and sporadically produce cohorts of meiotic primary spermatocytes [15,16]. In contrast to mitosis, homologous chromosomes are aligned during meiosis I, and the molecular machinery that facilitates interchromosomal HDR predominates over EJ [17]; either or both of these conditions might allow genotype conversion to occur in meiotic but not mitotic cells. We therefore interpreted our data to suggest that timing Cas9 expression to coincide with meiosis I may be critical to allow for genotype conversion in both sexes.

Here we use a single transgene to evaluate the importance of initiating Cas9 expression during prophase of meiosis I for genotype conversion to occur in mice. We engineered a knock-in allele at the *Spo11* locus to express Cas9 in conjunction with endogenous Spo11 using a P2A self-cleaving peptide (Fig 1A). Spo11 is the endonuclease and topoisomerase that catalyzes DSB formation required for crossing over during meiotic recombination in prophase of meiosis I [16,17]. For this analysis, the timing but not the catalytic activity of *Spo11* expression is important. In mammals, *Spo11* is expressed only in mature testes and in embryonic ovaries, and its expression peaks during prophase of meiosis I when homologous chromosomes are aligned [18,19].

**Fig 1.**
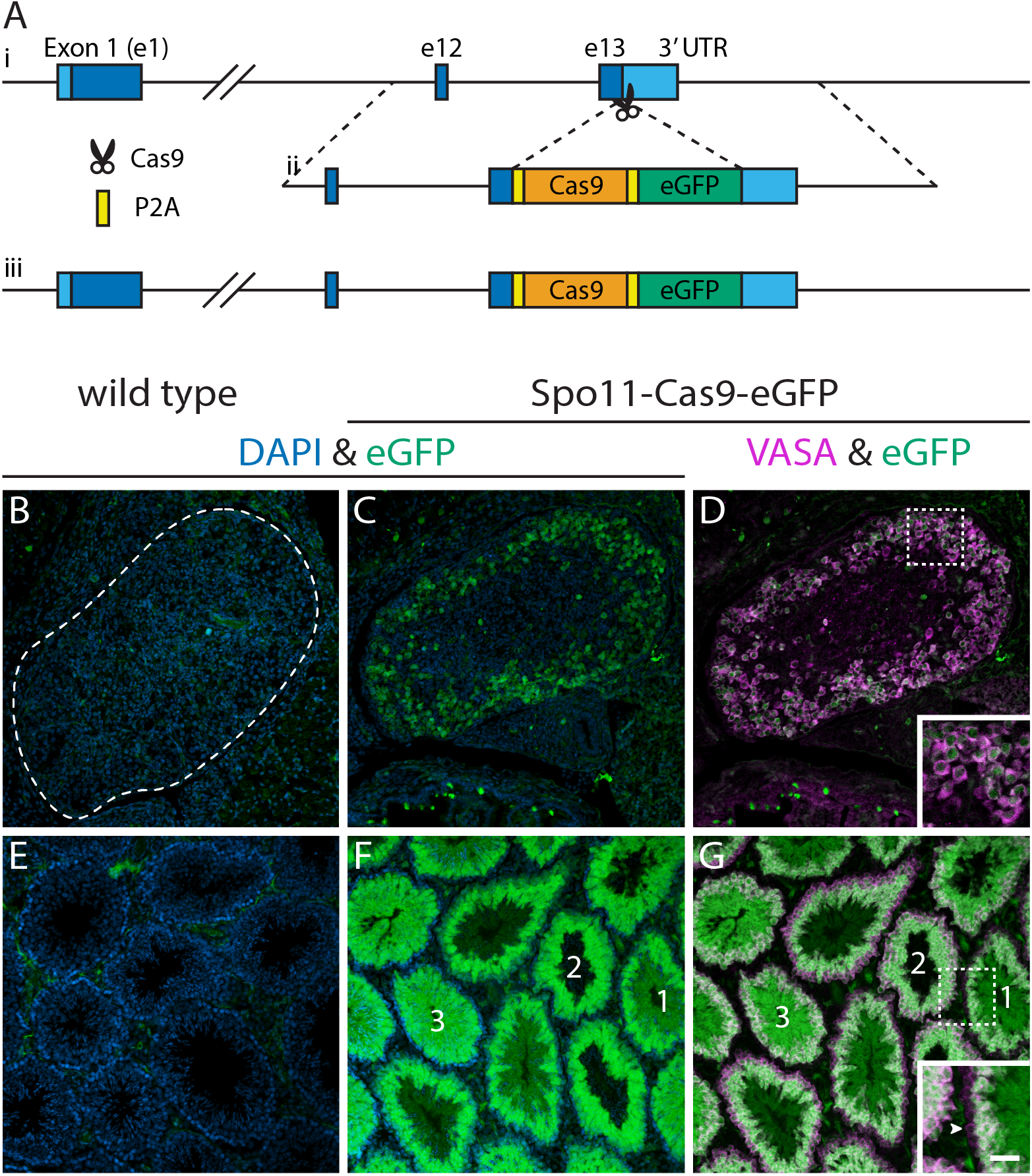
Male and female germ cell characterization of *Spo11*^*Cas9-P2A-eGFP*^ knock-in expression. **(A)** Schematic of targeting the *Spo11*^*Cas9-P2A-eGFP*^ knock-in allele. *i* Wild type *Spo11* locus. *ii* Construct for insertion. *iii* Final *Spo11*^*Cas9-P2A-eGFP*^ knock-in locus. The genotyping strategy is depicted in S1 Fig. **(B-G)** Immunofluorescence detection of eGFP (green) and VASA (magenta) plus DAPI (blue) in *Spo11*^*+/+*^ and *Spo11*^*Cas9-P2A-eGFP/+*^ mice. (B-D) embryonic ovaries at E18.5 (n=3) and (E-G) adult testes (n=3). Seminiferous tubules labeled 1 -> 3 are in different stages of maturation as determined by chromatin compaction (DAPI) and relative level of VASA expression in outer perimeter cells. The inset shows outer perimeter cells of tubule 1 (arrowhead points to spermatogonium) that express low VASA and no eGFP. By contrast, the outer perimeter cells of tubule 2 (meiotic primary spermatocytes) express low eGFP and higher VASA, and outer perimeter cells of tubule 3 express the highest eGFP. Scale bar: 50 µm for (B-G), 25 µm for the insets in (D) and (G). eGFP channel brightness was adjusted in (B-D) independent of (E-G); equivalent exposures are shown in S3 Fig.

Using the same *Tyr*^*CopyCat*^ allele and crossing scheme to assess genotype conversion efficiency as in our prior work [1], we demonstrate that Cas9 expression driven by *Spo11* during meiosis I promotes genotype conversion in the male as well as female germline. Compared to our most efficient prior strategy, the relative fraction of DSB repair that is mediated by HDR [(HDR/(HDR + EJ)] is increased in the male (from 0% to 11%) and is similar in the female (71% versus 67%) consistent with our hypothesis that restricting DSB repair to meiosis I favors HDR. However, total rates of DSB repair (HDR + EJ) are lower in both sexes resulting in absolute genotype conversion frequencies that are low in males and reduced in females compared to previous strategies. We suggest that the level of Cas9 expression driven from the *Spo11* locus may be too low for efficient DSB formation, though the timing of expression overlaps the window when interchromosomal HDR does occur. These results demonstrate that genotype conversion is possible in the male mouse germline and emphasize the importance of both timing and levels of Cas9 expression for efficient genotype conversion systems in rodents.

## Results

As summarized above, we hypothesized that restricting Cas9 expression to meiosis I in both the male and female germline should result in genotype conversion events in both sexes. *Spo11* expression is initiated in leptotene primary spermatocytes and oocytes and increases during zygotene [18,19]. In males, expression continues to rise through pachytene and diplotene and can be detected in round spermatids but not in residual bodies of elongating spermatids [18]. In females, expression falls through pachytene and diplotene, when oocytes arrest until ovulation [21]. We used a knock-in approach to place Cas9 after the final exon of *Spo11* to take advantage of the endogenous timing of expression from this locus while attempting to minimize disruption of *Spo11*, which is critical for fertility in mice. In this allele, Spo11, Cas9, and eGFP (to visually report expression from the locus) are each separated by self-cleaving peptide P2A sequences (Fig 1A). Hereafter, we refer to this transgene as *Spo11*^*Cas9-P2A-eGFP*^. Despite our efforts to preserve endogenous *Spo11* function, we recovered no offspring from a homozygous male after 20 months of continuous harem mating. Section histology of the testes revealed this homozygous *Spo11*^*Cas9-P2A-eGFP*^ male mouse had small testes and defects of the seminiferous tubule (S2 Fig) similar to *Spo11*^*-/-*^ mice [22].

We predicted that the regulatory context of the *Spo11* locus would limit Cas9-P2A-eGFP expression to meiotic spermatocytes and oocytes. We tested this hypothesis by performing immunofluorescence (IF) to detect transgene expression. Though we were unable to directly detect genetically-encoded Cas9 expression by IF or Western blot using commercially available Cas9 antibodies, we detected eGFP by IF in tissue sections of adult testes and female ovaries. In males, waves of spermatogenesis pulse along the length of seminiferous tubules such that a single cross section captures multiple phases [23,24]. Meiotic and maturing spermatocytes strongly express eGFP, which was not detected in cohorts of mitotically expanding spermatogonia (Fig 1E-G). In females, eGFP expression is first detected in ovaries at embryonic day 16.5 (E16.5), coinciding with early zygotene [21]. Expression increases in ovaries at E17.5 (late zygotene) and persists at E18.5. We note that the level of expression from the Spo11-Cas9-eGFP transgene appears to be higher in the male germline than in the female germline as determined by intensity of eGFP detection by immunofluorescence (Fig 1B-G and S3 Fig).

To assess the efficiency of genotype conversion using this transgene, we first crossed heterozygous *Spo11*^*Cas9-P2A-eGFP*^ knock-in mice with homozygous *Tyrosinase*^*chinchilla*^ (*Tyr*^*ch/ch*^) mice to introduce an ultra-tightly linked SNP in exon 5 of *Tyrosinase*. The *Tyr*^*ch*^ SNP is easily identified by PCR followed by Sanger sequencing and marks the recipient chromosome that will be targeted for DSB formation, allowing us to detect even a single genotype conversion event. We then crossed *Spo11*^*Cas9-P2A-eGFP/+*^*;Tyr*^*ch/ch*^ male mice to *Tyr*^*CopyCat/+*^ female mice (Fig 2B) to limit the possibility of maternal Cas9 transmission that might induce indel mutations in the early zygote [1].

**Fig 2.**
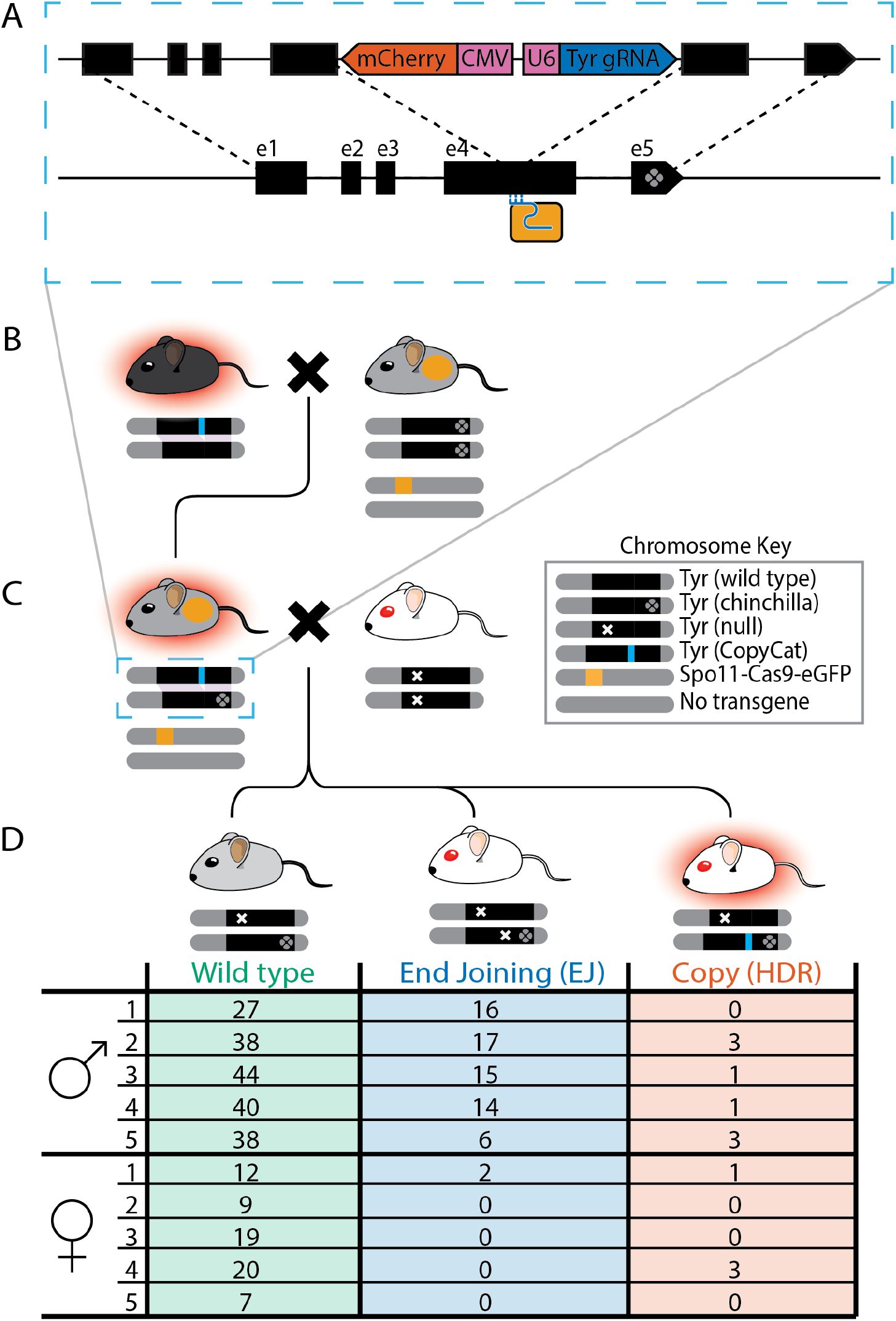
Breeding scheme to achieve genotype conversion with summary table of results in *Tyr*^*ch*^ offspring of complementation test crosses. **(A)** Schematic of the genetically encoded *Tyr*^*CopyCat*^ transgene in Exon 4 of *Tyrosinase* (top chromosome) and its insertion into the recipient chromosome (bottom) during interchromosomal HDR. mCherry (red), CMV and U6 promoters (pink), and Tyr4a-gRNA (dark blue). Dotted lines indicate *Tyr* homology between chromosomes. Cas9 (light orange box) and gRNA (light blue) only target the wild type *Tyr* allele. **(B**) Female *Tyr*^*CopyCat/+*^ crossed with male *Spo11*^*Cas9-P2A-eGFP/+*^*;Tyr*^*ch/ch*^ mice. **(C**) Complementation test cross of *Tyr*^*CopyCat/ch*^;*Spo11*^*Cas9-P2A-eGFP/+*^ male or female mice to albino *Tyr*^*null/null*^ mice. **(D)** Numbers of *Tyr*^*ch*^ offspring with each genotype in families (1-5) from male or female *Tyr*^*CopyCat/ch*^;*Spo11*^*Cas9-P2A-eGFP/+*^ parents. “Chromosome Key” shows genotypes of pictographic chromosomes under each mouse.

Cas9 expressed from this transgene in meiotic cells complexes with the Tyr4a-gRNA encoded by the *Tyr*^*CopyCat*^ transgene, which targets exon 4 of *Tyr* only on the *Tyr*^*ch*^-marked recipient chromosome (Fig 2A). A DSB may then be non-conservatively repaired by EJ to form an indel or by interchromosomal HDR to result in genotype conversion. If genotype conversion occurs, the *Tyr*^*CopyCat*^ transgene will be encoded in *cis* with the *Tyr*^*ch*^ SNP allele, which is expected to rarely occur between these loci by natural recombination, with a probability of 4.7 ⨯ 10^−5^.

To determine the frequency of these outcomes by genetic complementation, we crossed *Spo11*^*Cas9-P2A-eGFP/+*^; *Tyr*^*CopyCat/ch*^ mice to *Tyr*^*null/null*^ albino mice that have a null mutation in *Tyrosinase* exon 1 (Fig 2C). We genotyped all offspring of this cross for the *Tyr*^*ch*^ SNP (Fig 2D) and assessed coat color and mCherry fluorescence (S4 Fig) of the *Tyr*^*ch*^-positive population. Grey mice represent no cut or an in-frame repair at the gRNA target site in exon 4 that preserves activity of the *Tyr*^*ch*^ allele, while white mice have inherited a null mutation on the recipient *Tyr*^*ch*^ chromosome that fails to complement the *Tyr*^*null*^ allele. Of these, we identified white mice that also fluoresce red due to inheritance of the *Tyr*^*CopyCat*^ transgene that disrupts *Tyr* function (Fig 2D, right-most mouse). We then genotyped these red fluorescent *Tyr*^*ch*^ mice to confirm inheritance of the *Tyr*^*CopyCat*^ allele (S5 Fig). Since the *Tyr*^*null*^ mouse could contribute neither the *Tyr*^*ch*^ SNP nor the *Tyr*^*CopyCat*^ transgene, a mouse with both alleles must have inherited them on a single parental chromosome resulting from a germline interchromosomal HDR event.

We assessed the progeny of five males and five females using this complementation test cross strategy (Fig 2C and 2D). In total, we identified 263 *Tyr*^*ch*^-positive offspring of males and 73 *Tyr*^*ch*^-positive offspring of females. Of these, 8 offspring of males (3% of *Tyr*^*ch*^) and 4 offspring of females (6% of *Tyr*^*ch*^) inherited the *Tyr*^*CopyCat*^ transgene by genotype conversion. Although these numbers are far lower than the average 44% genotype conversion we previously reported in the female germline (*Vasa-Cre;H11-LSL-Cas9* strategy), this is the first reported observation of CRISPR/Cas9-mediated genotype conversion in the male germline.

Notably, Cas9 driven from the *Spo11* locus reduced the absolute rate of non-conservative DSB repair [(HDR + EJ)/total *Tyr*^*ch*^) compared to our previous *Vasa-Cre;H11-LSL-Cas9* conditional strategy from 100% to 29% in the male germline (Fig 3A), and from 60% to 8% in the female germline (Fig 3B). This reduction in DSB formation suggests the overall CRISPR-based cleavage efficiency is lower using the *Spo11*^*Cas9-P2A-eGFP*^ transgene than for previously employed strategies based on *Vasa*-mediated conditional Cas9 expression. We note, however, that the proportion of HDR events among all DSB repair outcomes [HDR/(HDR + EJ)] remains similar in females (71% in the previous study versus 67% in this study) and disproportionately improves in males (from 0% to 11%).

**Fig 3.**
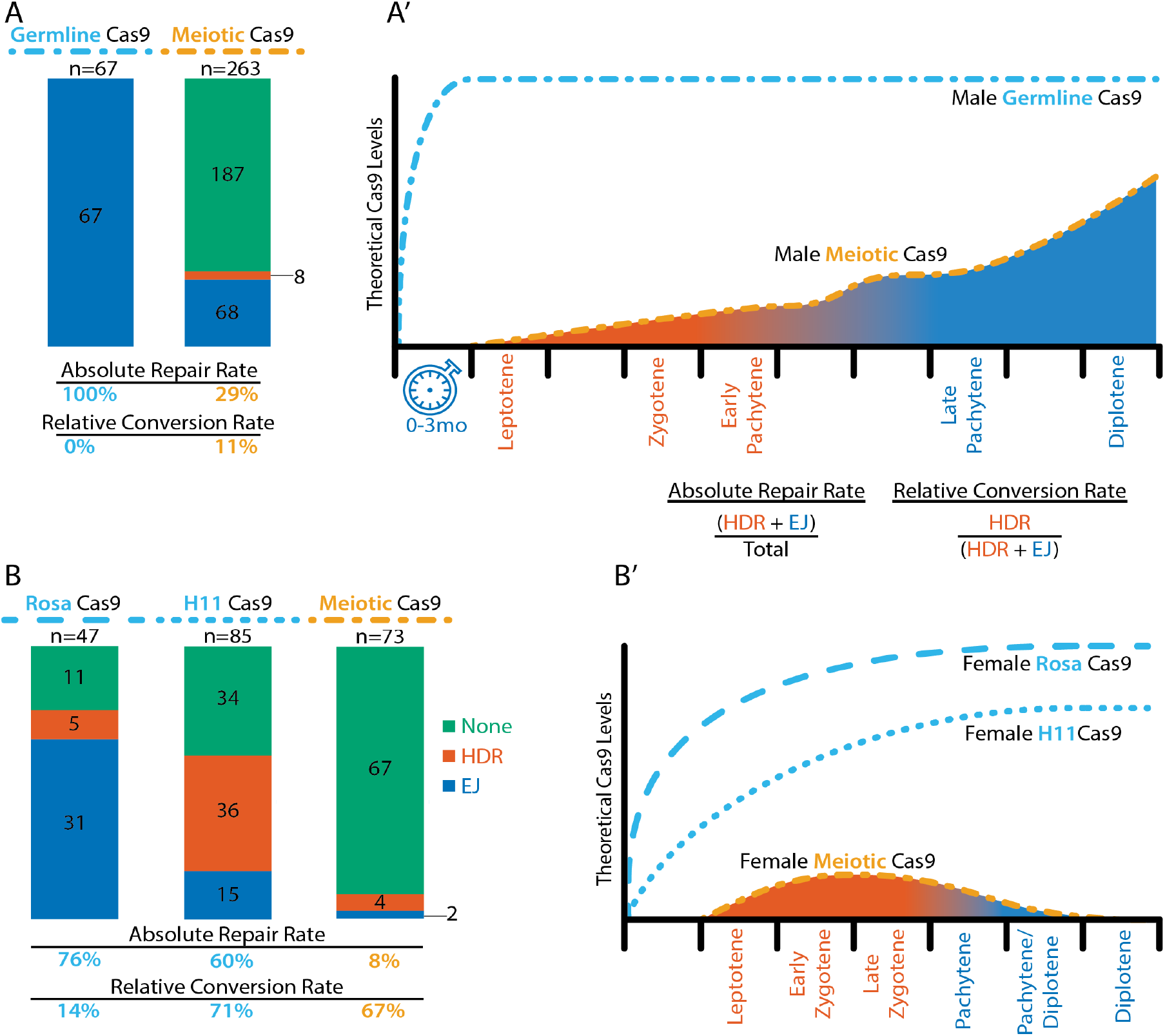
Comparisons of outcomes of each genetic strategy with models of theoretical Cas9 levels and timing with respect to DSB repair during meiosis I. **(A)** ‘Germline’ Cas9 offspring of males for the combined outcomes of Vasa-Cre;Rosa- and H11-LSLCas9, data sourced from [1], and ‘Meiotic’ Cas9 offspring of males from Fig 2D. **(B)** ‘Germline’ Cas9 offspring of females of each of the Vasa-Cre;Rosa- and H11-LSLCas9 ‘Germline’ strategies and ‘Meiotic’ Cas9 offspring of females from Fig 2D. Each bar graph represents the sum of repair types across all *Tyr*^*ch*^ recipient chromosomes in each strategy. Green, no cut or in frame repair; Red, interchromosomal homology directed repair (HDR); Blue, end joining, (EJ). Below each bar graph are the absolute repair rates and relative conversion rates, as defined by the equations shown. **(A’ and B’)** Models of theoretical Cas9 levels across the range of time relevant to genotype conversion. Each hash on the x-axis represents approximately one day. Leptotene in the female occurs at about E15.5 and initiates sporadically in maturing males. The orange line is the theorized relative Cas9 level inferred from RT-qPCR levels [21] and our eGFP IF ovaries and testes. The blue dotted line is theoretical expression of Cas9 under a strong CMV promoter and initiated in the germline by expression of Vasa-Cre in spermatogonia and oogonia. The blue dotted line plateaus faster in the male than in the female because many months are represented in the interval (clock) before meiosis is initiated in the male germline. The area under the curve and their respective stages are colored red (HDR) or blue (EJ) to demonstrate the inferred preference for the respective repair type at the time of the cut [17]. The relative shaded area of each color (and thus repair type) under each curve is roughly proportional to the observed “Relative Conversion Rate” shown below the respective ‘Meiosis’ bar graphs in (A) and (B).

## Discussion

Although moderately efficient genotype conversion in one parent might be sufficient to increase the probability of obtaining complex multi-locus genotypes, such approaches would be vastly improved by genotype conversion in both females and in males using a single Cas9 transgene. Direct expression from a single *Vasa-Cas9* transgene did not facilitate detectable genotype conversion, though the recipient chromosome was not marked [25]. Here, using a *Spo11*^*Cas9-P2A-eGFP*^ transgene, we have demonstrated the first reported genotype conversion in the male mouse germline. We also show that genotype conversion as a proportion of all repair events in the female germline is similar to our prior strategy. However, total repair events are substantially reduced in both sexes.

Together with our previous findings, our results suggest that the overall genotype conversion efficiency is dependent upon two key factors: the *absolute* rate of non-conservative DSB repair ([HDR + EJ]/total *Tyr*^*ch*^), and the *relative* rate of repair that is HDR (HDR/[HDR + EJ]) shown in Fig 3. Here, we synthesize all of our findings to suggest a model wherein these two rates are influenced by the cumulative amount of Cas9 expression and by the timing of Cas9 expression initiation. We suggest that both levels and timing of Cas9 dictate the dynamics of Cas9-mediated DSB formation and repair in context of the varied cellular and nuclear environments of male and female mitotic and meiotic germ cells.

The kinetics and mechanisms of DNA break repair have been historically described in the context of ionizing radiation and restriction enzyme-induced DSBs that are frequently repaired to preserve the original allele [26–31]. Recent work has shown that Cas9 induced DSBs may be repaired differently, as the Cas9 protein may remain bound after a DSB is induced and can thus prevent canonical repair machinery from accessing exposed DSBs [32– 34], though local chromatin environments can affect Cas9 binding kinetics [35,36]. These mechanics may explain observations that Cas9-induced DSB repair is substantially more error prone than repair after other means of DSB induction [34]. This suggests that non-conservatively repaired alleles are a reasonable proxy for total rate of DSB generation, as most conserved alleles in our crosses were likely never cut as opposed to cut and correctly repaired.

These studies inform our interpretation of observed differences in the absolute rate of DSB repair in the context of genetically encoded Cas9. In our previous strategies, after Vasa-Cre mediated removal of the lox-STOP-lox, Cas9 was expressed under regulatory control of the strong constitutive CAG promoter at either of the permissive H11 or Rosa26 loci. These strategies generated a non-conservative allele in 100% of *Tyr*^*ch*^ male germ cells (Fig 3A, all EJ) and in 60-76% of female germ cells [1] (Fig 3B, HDR + EJ). In contrast, the absolute rate of DSB repair in this meiotic *Spo11*^*Cas9-P2A-eGFP*^ strategy is substantially lower at 29% (76/263) in male (Fig 3A) and 8% (6/73) in female meiotic germ cells (Fig 3B). Previous studies demonstrated that Cas9-induced DSBs follow the dynamics of mass action; across a fixed window of time, total DSBs increase proportionally to the amount of Cas9 [37], and a higher proportion of DSBs occur earlier in the presence of higher levels of Cas9 [38]. Thus, we hypothesize that the lower absolute rate of DSB repair in the *Spo11*^*Cas9-P2A-eGFP*^ strategy may reflect lower Cas9 expression from endogenous *Spo11* regulatory sequences than from the constitutive CAG promoter at a permissive locus. We note that the difference in absolute rate of DSB formation in males and females of the *Spo11*^*Cas9-P2A-eGFP*^ strategy correlates with a higher level of eGFP detected in male meiotic germ cells (S3 Fig). This observed difference is also consistent with previously reported quantification of *Spo11* mRNA expression by RT-qPCR, which shows a higher expression level in males than in females [21].

It is essential to note that the total level of Cas9 protein in a cell is dependent not only on the strength of a regulatory sequence, but also on the duration of expression. The 100% absolute rate of DSB repair in the Vasa-Cre;LSL-Cas9 male germline strategies may have resulted from persistent Cas9 expression over several months in the population of self-renewing mitotic spermatogonia (Fig 3A’, blue clock). Such a long duration of Cas9 expression would provide ample time for prevalent DSB repair by EJ; resulting indels would be inherited by primary spermatocytes, thus blocking any subsequent DSB formation and genotype conversion at the onset of meiosis. By contrast, the *Spo11*^*Cas9-P2A-eGFP*^ transgene delays Cas9 expression by months to be initiated first in meiotic primary spermatocytes (Fig 1G and Fig 3A’, orange line), and expression termination is controlled by the endogenous *Spo11* locus. After rising during meiosis I, *Spo11* levels drop precipitously in maturing spermatids [21].

Together, this suggests that the absolute rate of DSB formation correlates with a cumulative level of Cas9 expression that is likely lower in the *Spo11*^*Cas9-P2A-eGFP*^ strategy than in the Vasa-Cre;LSL-Cas9 strategies and lower in *Spo11*^*Cas9-P2A-eGFP*^ females than in males.

In contrast to the absolute rate of DSB repair, we think that the relative rate of HDR [HDR/(HDR + EJ)] is primarily a function of the timing of Cas9 activity rather than the amount of Cas9 protein. In meiotic cells, particularly in early prophase I (leptotene through mid-pachytene), EJ-specific machinery is not detected; homologous recombination becomes the exclusive repair pathway and predominantly occurs between homologous chromosomes [39,40]. Later in prophase I, after mid-pachytene, EJ again becomes the primary DSB repair choice [17]. Second, the single homologous chromosome that is the template for genotype conversion is far more likely to be aligned and accessible during meiosis, and machinery is expressed that limits recombination to occur between homologous chromosomes [17,41]. From these data, we hypothesize that a permissive window for genotype conversion by interchromosomal HDR opens in leptotene and closes in mid-pachytene of meiosis I (Fig 3A’ and 3B’, orange shading), whereas DSBs induced before or after this window may be more likely repaired by EJ.

The similar relative rate of HDR in females of both genetic strategies may reflect a narrow temporal difference between the onset of Cas9 expression in the *Spo11*^*Cas9-P2A-eGFP*^ strategy compared to the Vasa-Cre;LSL-Cas9 strategies, since Vasa-Cre is expressed in oogonia that rapidly mature into meiotic oocytes. By contrast, in the male Vasa-Cre;LSL-Cas9 strategies, DSBs may have been formed and repaired by EJ in most or all mitotic spermatogonia prior to opening the permissive window. Using the *Spo11*^*Cas9-P2A-eGFP*^ that initiates expression in meiotic primary spermatocytes brings the onset of Cas9 expression closer to the permissive window. Therefore, temporally shifting Cas9 to the onset of meiosis in both sexes resulted in disproportionate improvement of relative rate of HDR in males versus females.

Our results mark a significant advance in the feasibility of super-Mendelian inheritance strategies in rodents by achieving germline genotype conversion in both sexes. We propose that adjusting the onset and increasing the level of Cas9 expression can further improve the absolute rate of DSB repair and the proportion of repair that is genotype conversion by HDR in both males and females. This may be achieved by identifying new promoters that express at a high level and that fall within or immediately prior to the permissive window for HDR during meiosis I. An alternative approach might be to use a promoter that drives strong expression with less temporal specificity and to chemically control the timing of Cas9 activity (e.g. with tamoxifen or trimethoprim [42]). Both wild release applications to constitute a ‘gene drive’ and laboratory applications to facilitate breeding complex genotypes will benefit from further increasing the efficiency of genotype conversion by improved genetic strategies. However, improvements that require drug delivery would not be practical for implementation in the wild.

Increasing the absolute rate of DSB repair versus the relative rate of HDR will have different effects on improving gene drive versus laboratory applications. Gene drive efficiency can tolerate a lower absolute rate of repair as long as the proportion of repair that is HDR is very high. Otherwise systems must be designed to select against indels that would be resistant to subsequent genotype conversion (reviewed in [43]). In a laboratory setting, where the researcher can genotype to choose animals, a higher rate of EJ is more tolerable so long as the rate of HDR is sufficient to obtain genotypes of interest. While showing that CRISPR/Cas9-mediated genotype conversion is possible in both male and female mouse germlines, our work reveals nuances that differ from insects and that must be considered for further refinement and implementation in rodents.

## Acknowledgements

We thank H. Cook-Andersen, M. Wilkinson, and R. Knight for discussions of strategy and meiotic drivers. C. Weber and V. Ruthig provided advice on germline immunofluorescence and staging of ovaries and testes. We thank D. Yelon for use of the Leica TCS SP8 multiphoton confocal microscope. This work was funded by a Packard Fellowship in Science and Engineering from the David and Lucile Packard Foundation and NIH grant R21GM129448 awarded to K.L.C., NIH grant R01AI131081 awarded to S.M.H., an Allen Frontiers Group Distinguished Investigators Award to E.B., and a gift from the Tata Trusts in India to TIGS-UCSD and TIGS-India. A.J.W. and H.A.G. were supported by NIH training grant T32GM007240, and R.L. was supported by NIH training grant T32GM133351.

## Methods

### Animals

Production of the *Tyr*^*CopyCat*^ transgenic mice was previously described in [1]. *Tyr*^*chinchilla*^ mice are strain FVB.129P2-Pde6b+ Tyr^c-ch/AntJ^, stock number 4828, from the Jackson Laboratory. *Tyr*^*null*^ mice are strain CD-1, stock number 022, from Charles River Laboratories. The *Spo11*^*Cas9-P2A-eGFP*^ knock in mice were produced by Biocytogen using CRISPR/Cas9 to insert P2A-Cas9-P2A-GFP flanked by 1.5 kb arms of homology to *Spo11* exon 13 and the 3’ UTR. All mice were housed in accordance with UCSD Institutional Animal Care and Use Committee protocols and fed on a standard breeders diet.

### Sample Collection, Immunofluorescence, Histology, and Imaging

We collected testes from adult male *Spo11*^*Cas9-P2A-eGFP/+*^ and wild type mice. To obtain meiotic ovaries, we crossed *Spo11*^*Cas9-P2A-eGFP/+*^ males to CD-1 females. Once fertilization was inferred by detection of a copulatory plug, the female was separated from the breeding male. At the desired embryonic stage, we dissected embryos from the uterus, confirmed the embryonic stage by limb morphological stage [44], bisected the embryo in transverse anterior to the ovaries, and collected the tail tip for PCR genotyping to establish sex and presence of the *Spo11*^*Cas9-P2A-eGFP*^ transgene. Embryos were fixed in 4% PFA overnight. The following day they were transferred to 20% sucrose solution until saturated. Embryos were embedded in OCT blocks, and sectioned at 6 µm thickness. Slides were frozen at -80°C until processing.

Slides were removed from -80°C and dried. We performed antigen retrieval for 10’ in 5µg/ml Proteinase K at room temperature. We then fixed in 4%PFA 1xPBS for 5’ and washed 3x for 10’ in 1x PBS. Slides were then blocked (3% BSA, 5% NGS, 0.1% Triton, 0.02% SDS, in 1x PBS) for 1-2 hr at room temperature before addition of primary antibodies diluted in block [chicken anti-eGFP, 1:500, Thermo Fisher catalogue # A10262, and rabbit anti-human Ddx4 (VASA), 1:250, Abcam catalogue # AB13840] and incubated at 4°C for 18 hours. Slides were then washed 3x for 30’ in 1x PBS; 0.1% Triton (PBST) prior to incubation in secondary antibodies [Invitrogen Alexa-488 goat anti-chicken and Life Technologies 647 goat anti-rabbit, 1:250 each] in block with 300 nM DAPI [Cell Signaling, catalogue # 4083S] and incubated at 4°C for 18 hours. We then washed 3x for 30’ in PBST and mounted under coverslips in Vectashield (Vector Labs).

Images were collected on Leica TCS SP8 multiphoton confocal microscope. Excitation frequency GFP (488 nm): emission range 505 – 550 nm. Excitation frequency Vasa (647 nm): emission range 655 – 685 nm. Excitation frequency DAPI (405 nm): emission range 415 – 470 nm.

The pixel intensity was determined in FIJI by ZProjection ‘average intensity’ for each channel. The brightness of eGFP signal intensity was adjusted equivalently across all samples (Fig 1 and S3 Fig) with the exception of the testis in Fig 1F and 1G, which was reduced for clarity.

For hematoxylin and eosin staining of slides, the staining agent was Harris hematoxylin solution. Samples were differentiated in 1% acid alcohol. The bluing agent was saturated lithium carbonate and slides were counterstained in a solution of 0.1% eosin Y and 0.1% phloxine B in acid alcohol. The samples were then dehydrated to 100% EtOH and cleared in xylenes. The slides were mounted under coverslips in Permount and cured for 24 hours. Samples were imaged at 20X magnification under brightfield on an Olympus BX61 microscope.

### Genotyping

The *Spo11*^*Cas9-P2A-eGFP*^ knock in allele was detected as in S1 Fig using primers and conditions in S1 Table. We detected inheritance of the *Tyr*^*CopyCat*^ transgene by observing red fluorescence of mCherry (S4 Fig). We then genotyped all animals for the *Tyr*^*ch*^ allele by PCR amplification of a 392 bp region containing the SNP followed by Sanger Sequencing (GeneWiz). Finally, we confirmed the *Tyr*^*CopyCat*^ genotype in each *Tyr*^*ch*^ animal using PCR primers that bind inside and outside of the transgene to amplify an 838 bp fragment only from transgenic animals (S5 Fig). PCR genotypes were performed according to conditions set in S1 Table.

## Supporting Information

**S1. Fig.**
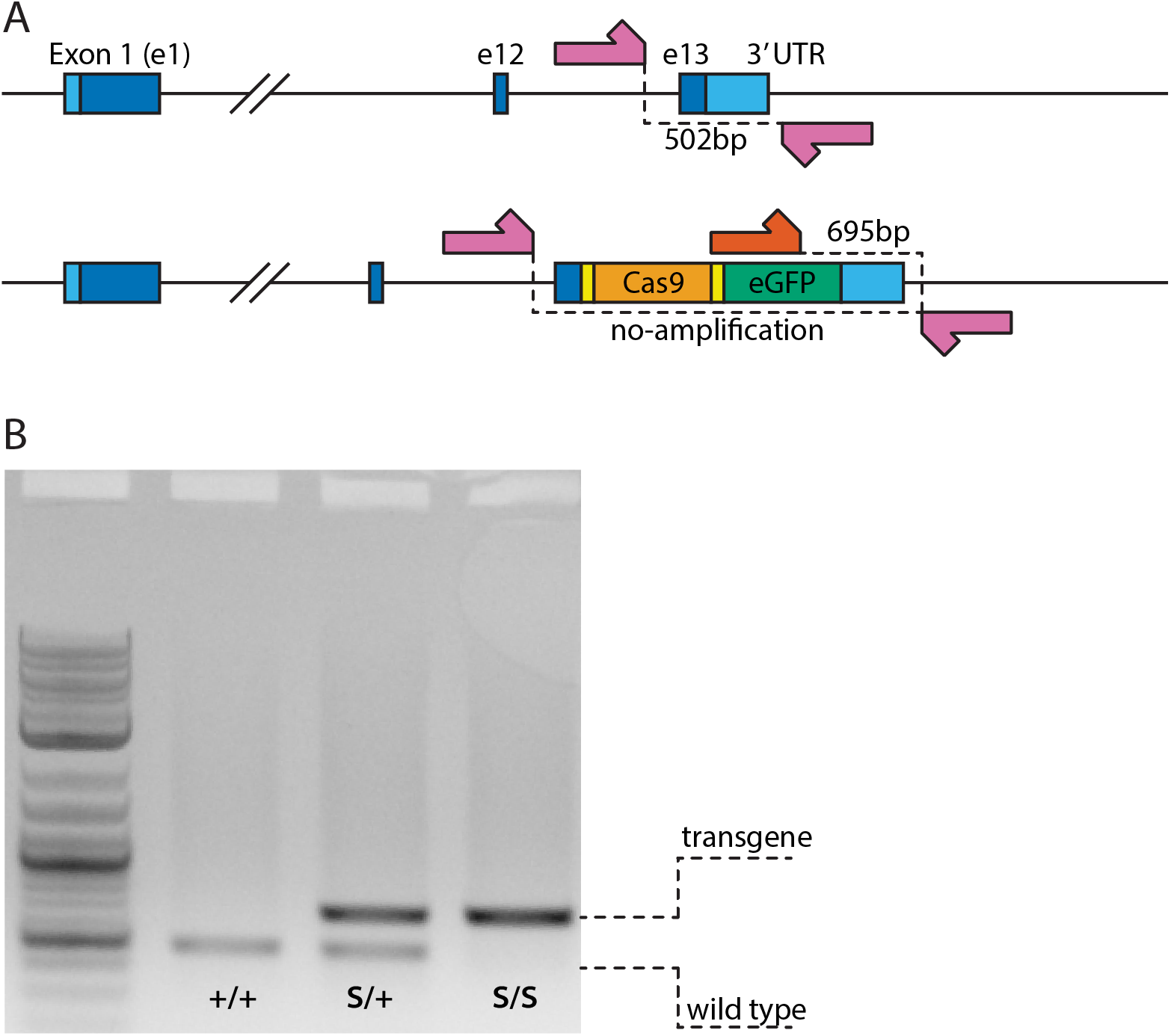
Genotyping strategy for *Spo11*^*Cas9-P2A-eGFP*^. **(A)** Schematic of primer binding location for PCR genotyping. **(B)** Gel depicting PCR of Spo11 locus revealing the genotype of *Spo11^+/+^*, *Spo11^Cas9-P2A-eGFP/+^*, and *Spo11^Cas9-P2A-eGFP/Cas9-P2A-eGFP^*.

**S2. Fig.**
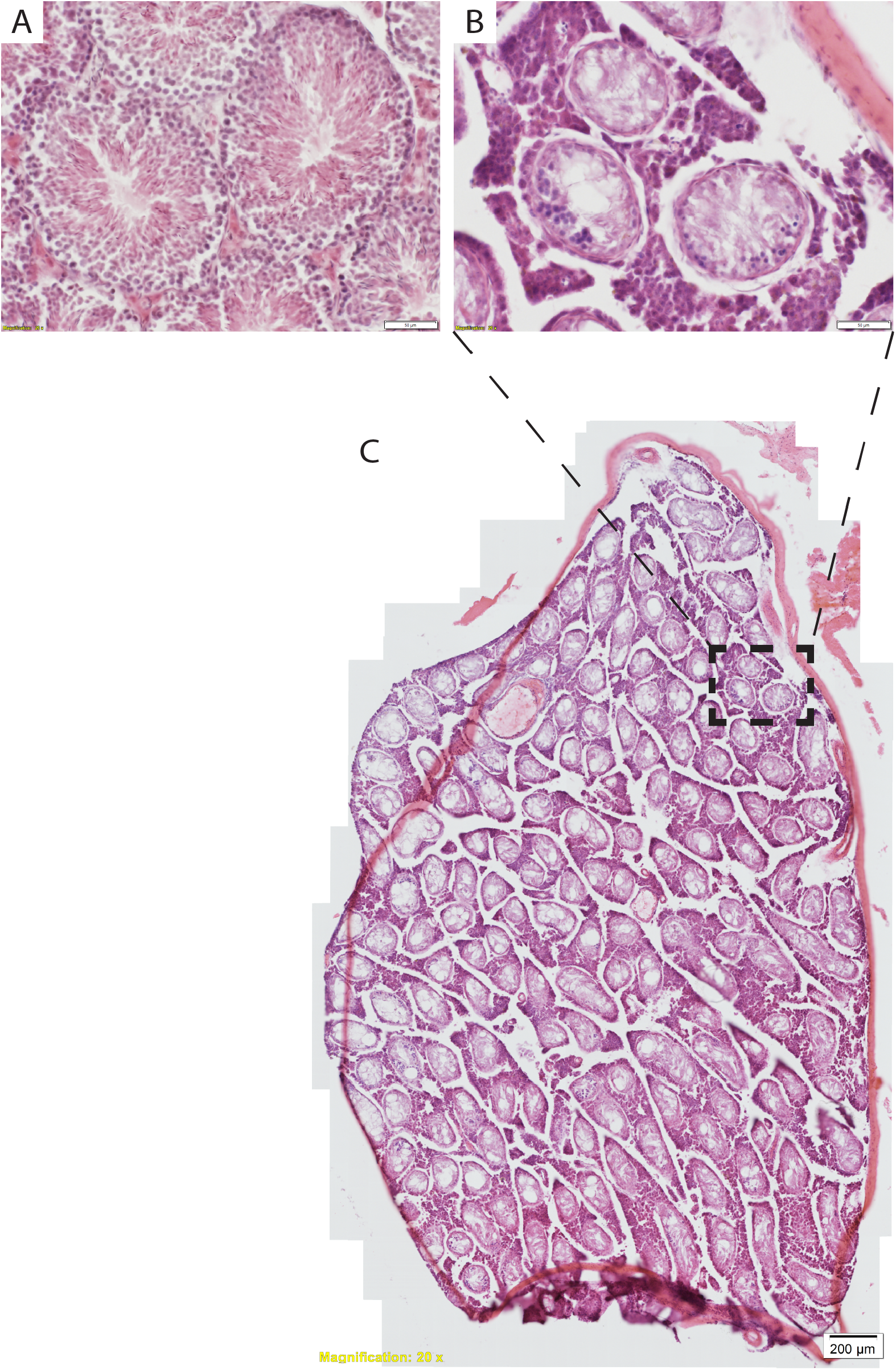
Hematoxylin and eosin staining of testes. **(A)** *Spo11*^*Cas9-P2A-eGFP*/*+*^ and **(B)** *Spo11*^*Cas9-P2A-eGFP*/*Cas9-P2A-eGFP*^ seminiferous tubules; scale bar is 50 µm. Cells in seminiferous tubules in (B) likely do not complete meiosis, evidenced by absence of sperm. **(C**) Tiled image of *Spo11*^*Cas9-P2A-eGFP*/*Cas9-P2A-eGFP*^ testis; scale bar is 200 µm. All seminiferous tubules are deformed and devoid of sperm.

**S3. Fig.**
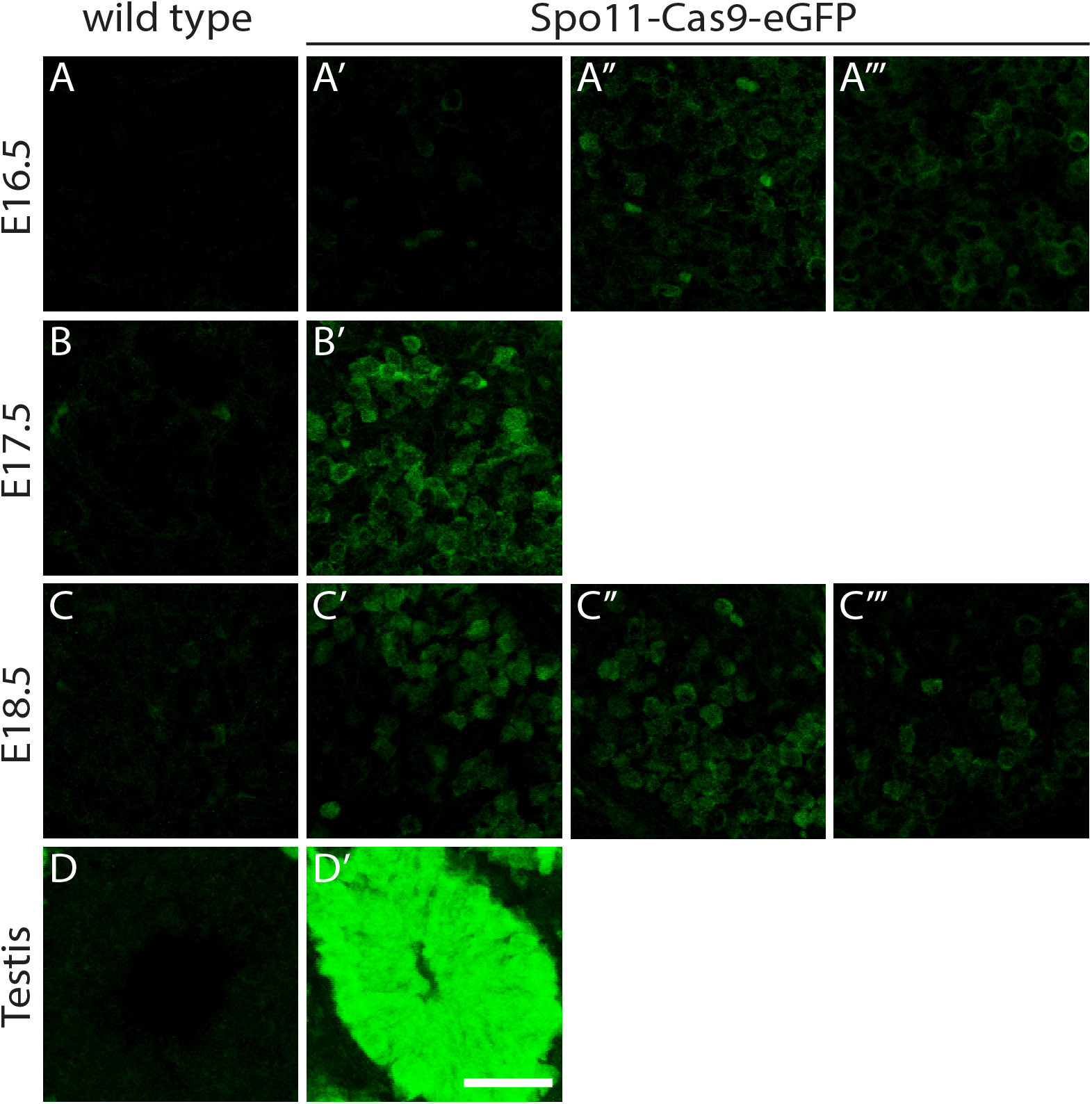
Immunofluorescence detection of eGFP in embryonic ovaries and mature testis. **(A-C)** Wild type embryonic ovary. All other panels in each row are *Spo11*^*Cas9-P2A-eGFP*/+^ littermates (e.g. A’-A’’’). **(D)** Wild type and **(D’)** *Spo11*^*Cas9-P2A-eGFP*/+^ adult testis. Scale bar is 50 µm for all panels. Confocal fluorescence imaging settings are identical for all samples; eGFP expression is substantially higher in testes compared to any embryonic ovary time point

**S4. Fig.**
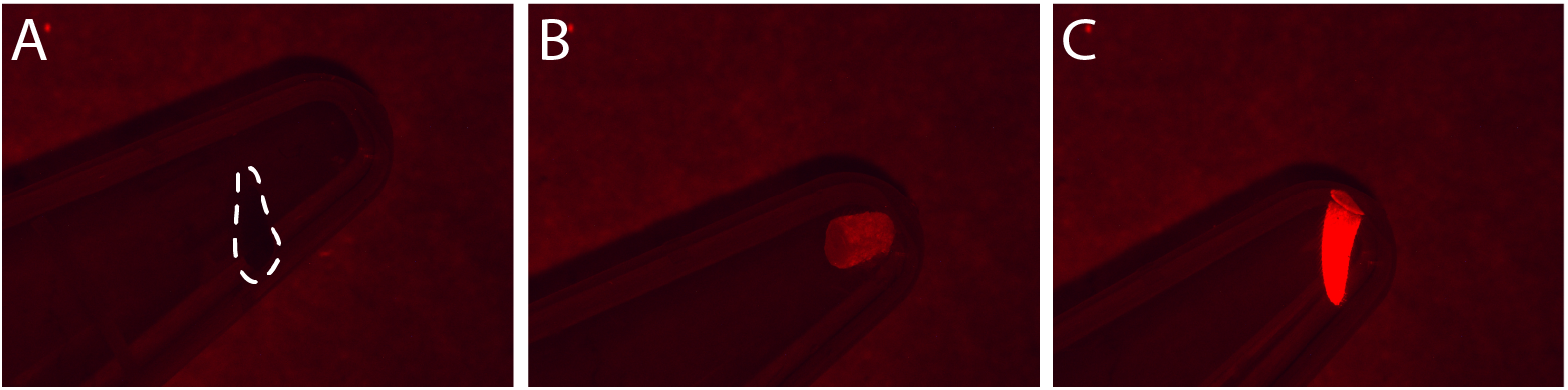
Detection of CopyCat transgene by red flluorescent mCherry in tail tips. **(A)** *Tyr*^*null*^*/Tyr*^*null*^ tail tip. **(B-C)** *Tyr*^*CopyCat*^*/Tyr*^*null*^.tail tips. While clearly present or absent, mCherry expression varies in intensity between individuals of equivalent age.

**S5. Fig.**
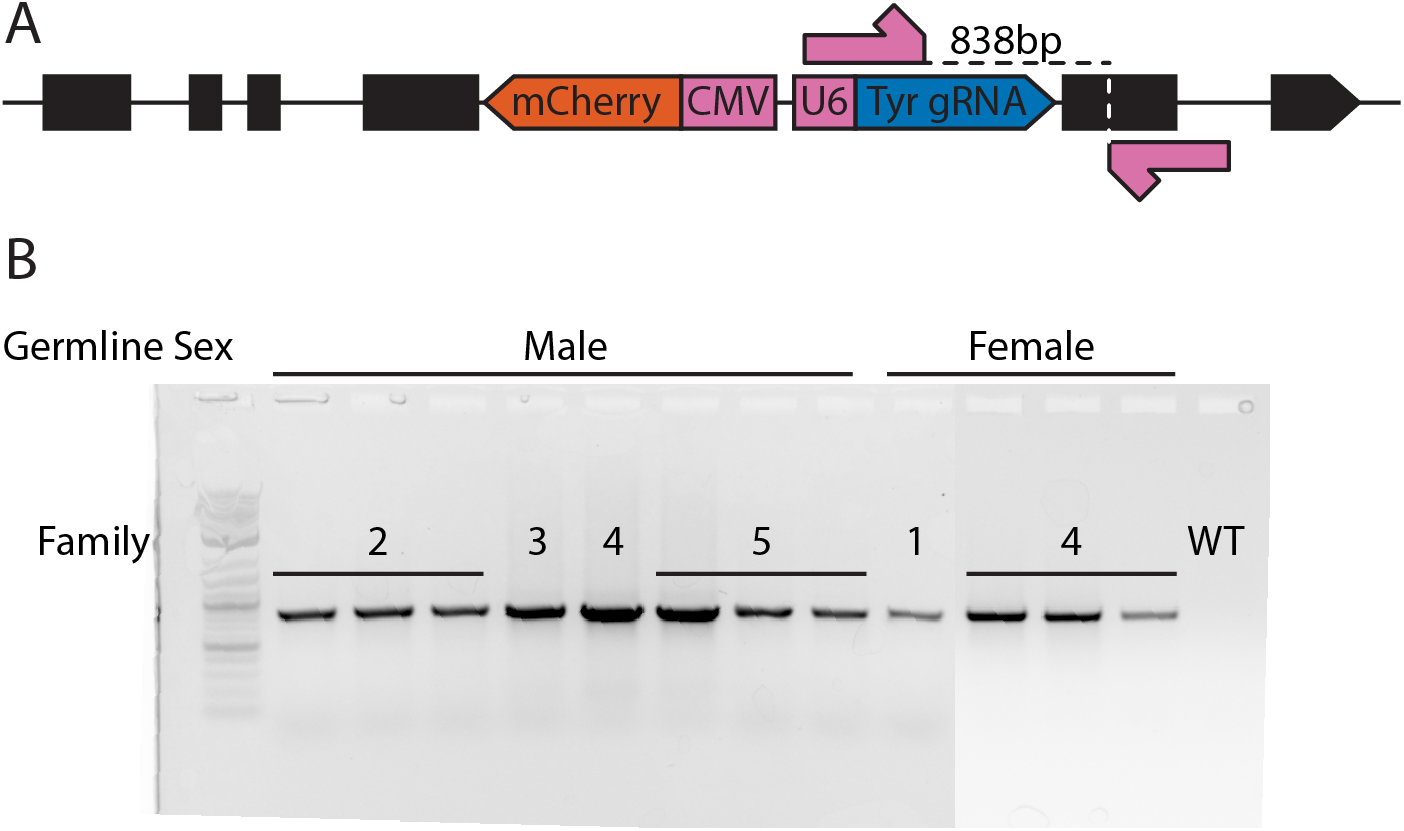
PCR confirmation of *Tyr*^*CopyCat*^ in each offspring of an HDR event. **(A)** Schematic of primer binding locations to detect presence of the *Tyr*^*CopyCat*^ transgene. **(B)** PCR confirmed presence of the *Tyr*^*CopyCat*^ transgene in each of the individuals marked as positive for a genotype conversion event, with families of offspring from males and females numbered as in the table in Fig 2D. Two gels were merged for ease of understanding.

**S1. Table.**
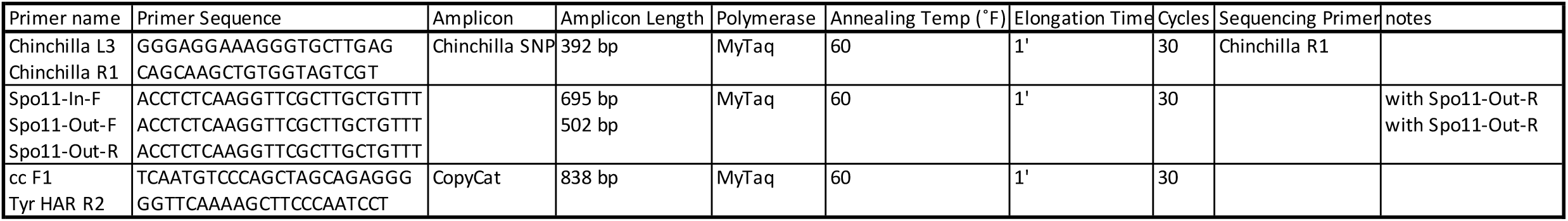
Primer sequences and PCR conditions for each genotyping strategy.

